# Spectrogram cross-correlation can be used to measure the complexity of bird vocalizations

**DOI:** 10.1101/2021.03.30.437665

**Authors:** Suyash Sawant, Chiti Arvind, Viral Joshi, V. V. Robin

## Abstract

1. Birdsong is an important signal in mate attraction and territorial defense. Quantifying the complexity of these songs can shed light on individual fitness, sexual selection, and behavior. Several techniques have been used to quantify song complexity and be broadly categorized into diversity indices, measures of stationary probabilities, and measures of sequential variations. However, these methods are unable to account for important acoustic features like the frequency bandwidth and the variety in the shape of syllables which are an integral part of these vocal signals. This study proposes a new complexity measure that considers intra-song note variability and calculates a weighted index for birdsongs using spectral cross-correlation.
2. We compared the previously described methods to understand the advantages and limitations based on the factors that would be affecting the complexity of songs. We developed a new method- Note Variability Index (NVI), which incorporates the spectral features of notes while quantifying complexity. This measure alleviates the need for manual annotations of notes that can be error-prone. We used Spectrogram Cross-Correlation (SPCC) to compare notes within a song and used the output values to quantify song complexity.
3. To check for the efficacy of the new method, we generated synthetic songs to caricature extremes in song complexity and compared selected conventional complexity measures along with the NVI. We provide case-specific limitations of these methods. Additionally, to examine the efficacy of this new method in real-world scenarios, we used natural birdsongs from multiple species across the globe with varying song structures to compare conventional methods with NVI.
4. To our knowledge, NVI is the only song complexity method that captures the variation of spectral features of notes in songs where the conventional methods fail to distinguish between similar song structures with different note types. As NVI does not need a manual classification of notes, it can be easily implemented for any type of birdsong with existing sound analysis softwares; it is very quick, avoids the possible biases in note classification, and can possibly be automated for large datasets in the future.

## Introduction

Vocal signals form an integral part of animal communication. The diversity and variability in these vocalizations serve several functional roles in foraging (Boogert, Giraldeau, & Lefebvre, 2008), attracting a mate (Byers & Kroodsma, 2009), and warning against predators (Demartsev et al., 2014). Each type of vocalization is unique and usually consists of a sequential order of notes, which vary in structure (frequency, duration, energy, pace, and intensity) and the number of repetitions (Marler & Slabbekoorn, 2004). Variation in notes gives rise to complexity, which is common in the songs of Passeriform birds and is of interest to ecologists and evolutionary biologists.

The complexity of birdsongs is often studied in two ways; a) Immediate Complexity- the complexity in vocalization within individual song bouts and b) Eventual Complexity- the complexity of vocalization across songs (László Z. Garamszegi et al., 2005). Birdsong complexity has often been attributed to a large repertoire size, resulting from open-ended learning (Robinson, Snyder, & Creanza, 2019), and mimicry (László Zsolt Garamszegi, Eens, Pavlova, Avilés, & Møller, 2007). Song complexity has been used to measure individual fitness, particularly in mate attraction and territorial defense (Catchpole, 1987; Horn & Bruce Falls, 2010). In many cases, birds having a more complex song have a higher reproductive success (Nowicki, Peters, & Podos, 1998; Cramer, 2013). While various ecological and evolutionary studies rely on understanding the relationship between song complexity and sexual selection, quantifying song complexity has remained a challenge with no uniform measure.

In this study, the term ‘song’ is used for a series of notes and syllables separated by visible time gaps. A note is the smallest unit in a song and can be defined in various ways described in Kershenbaum et al. (2016).

Previous studies have measured birdsong complexity based on diversity, repetition, order, combinations of notes, and their delivery rate (Kershenbaum et al., 2016). These indices can be broadly classified into three categories-measures of the diversity of notes, measures of stationary probabilities (rarity) of notes, and measures of sequential variations of notes (order) within a song.

### Stationary probability (the rarity of note appearances)

A well-established ecological method, Shannon’s Diversity uses the relative abundance by measuring the stationary probabilities of notes to quantify song complexity (Kershenbaum, 2014). Two methods, Repeat Distribution and Mutual Information (Kershenbaum & Garland, 2015) also used the stationary probabilities of notes within a song. These measures incorporate the occurrence of rare notes as higher complexity but don’t consider the range of spectral properties of songs; additionally, this method also needs notes to be manually classified into distinct types (See Supp. Fig 1 for an illustration).

### Order of notes

To understand the variations in the order of notes, songs are annotated as sequences of distinct elements. Markov models have been commonly used to understand transition probabilities of notes within a song (Katahira, Suzuki, Kagawa, & Okanoya, 2013; Kershenbaum & Garland, 2015). Other methods that consider the order of notes include Lempel-Ziv Complexity, Levenshtein Distances, and Entropy rate (Kershenbaum, 2014; Kershenbaum & Garland, 2015). (Sasahara, Cody, Cohen, & Taylor, 2012) used Song Directed Networks and Song Undirected Networks to analyze the order of notes within a song which utilizes deterministic and non-deterministic transition motifs to quantify the complexity of songs. These measures help in understanding the variation in the order of notes (as pre-defined by the researchers) but may not always account for the song complexity in terms of structural diversity of notes as no spectral parameters are considered.

### Diversity of notes

Many studies (László Z. Garamszegi et al., 2005; Zeng et al., 2007; Petrusková, Pišvejcová, Kinštová, Brinke, & Petrusek, 2016; Benedict & Najar, 2019) used repertoire size (by characterizing note types) to understand the overall note diversity for a species. Although repertoire size captures the eventual note variety, it may not represent the complexity within a song as a bird with a larger repertoire size may repeat the same notes within a song (low immediate variety) (e.g., Brown Thrasher), and a bird with a limited repertoire may create complex songs from combinations of very few notes (e.g., Gray-headed canary-Flycatcher). Other commonly used metrics of the diversity of notes include simple metrics such as the number of notes within a song, the types of notes within a song, or the ratio of the type of notes and number of notes (Spencer, Buchanan, Goldsmith, & Catchpole, 2003). Song Variability Index used variation in song structure to compare the diversity across songs (Singh & Price, 2015; Purushotham & Robin, 2016).

Along with the complexity measures mentioned above, some studies have also used specific acoustic parameters of songs to understand the variation (Benedict & Najar, 2019). Mason, Shultz, & Burns (2014) used eigenvalues from Principal Component Analysis of multiple acoustic features to study the complexity of songs. However, these methods are very case-specific as the results solely depend upon selecting song features chosen based on the signal form and delivery patterns of the study species (Kershenbaum et al., 2018).

More recently, Zsebők et al. (2021) used conventional ecological functional diversity metrics Functional Richness (Fric), Function Evenness (Feve), and Function Divergence (Fdiv) to estimate song complexity. The raw data for these methods also come from manual spectral measurements of maximum, minimum frequency, and time of each note. However, multiple note types with varying spectral shapes can have similar frequency and time bounds, and these may be missed by such methods.

The methods used so far have shown promising results in understanding birdsong complexity; however, there are a few major challenges. In measures that use note types, notes are always considered as discrete acoustic unit types. There can be a gradient in the variation in notes in real cases, and because of this, it is challenging and, in some cases, may even be biologically inappropriate to classify the notes into discrete categories manually. Manual classification can also be highly biased as it is based on aural and visual classification of the spectrogram. Other measures that use spectral parameters use only the manually ascertained minimum and maximum values (frequency and time) of a note, and such measures may miss the complete structure of a note, such as the shape of a note. Thus, the complexity of bird vocalizations can be measured as both diversity of notes within manually defined boundaries, using various analytical tools, but these have limitations of missing various spectral features and that of scalability (automation) with large datasets.

Here we describe a new complexity measure, Note Variability Index (NVI), that incorporates the weighted inter-note diversity within a song calculated using Spectrogram Cross-Correlation (SPCC) (Charif, Waack, & Strickman, 2010).

We compare the efficacy of our new metric, NVI, with two song complexity indices-Shannon’s equitability and note type by note count ratio using a) synthetic songs with different combinations of simulated notes to create a dataset with extremes, and b) natural bird songs from 12 global bird species that have vocalizations ranging from simple to complex. Within natural songs, we looked at both immediate and overall (eventual) note variety.

## Materials and Methods

### 2.1 Note Variability Index (NVI)

#### Describing NVI

For a song with N_C_ number of notes, the normalized correlation coefficients C_Δt_ ∈ (0,1) are obtained by evaluating correlation for every pair of notes, which forms an N_C_ x N_C_ similarity matrix. C_Δt_ values tending towards 1 represent a higher association between two notes, while C_Δt_ = 0 implies no association between the notes. (Fig. 1b). NVI is the cumulative score of the inverse correlation coefficients in the similarity matrix (1 - C_Δt_), where i and j are the row and column ranks. The denominator N_c_(N_c_ - 1) is used to obtain a normalized value to compare the song complexities across songs with an unequal number of notes.

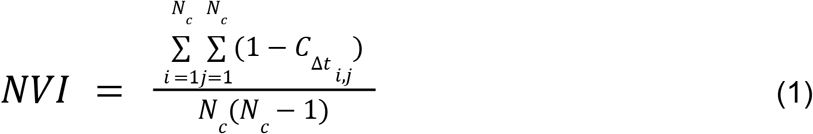

**Fig. 1.**
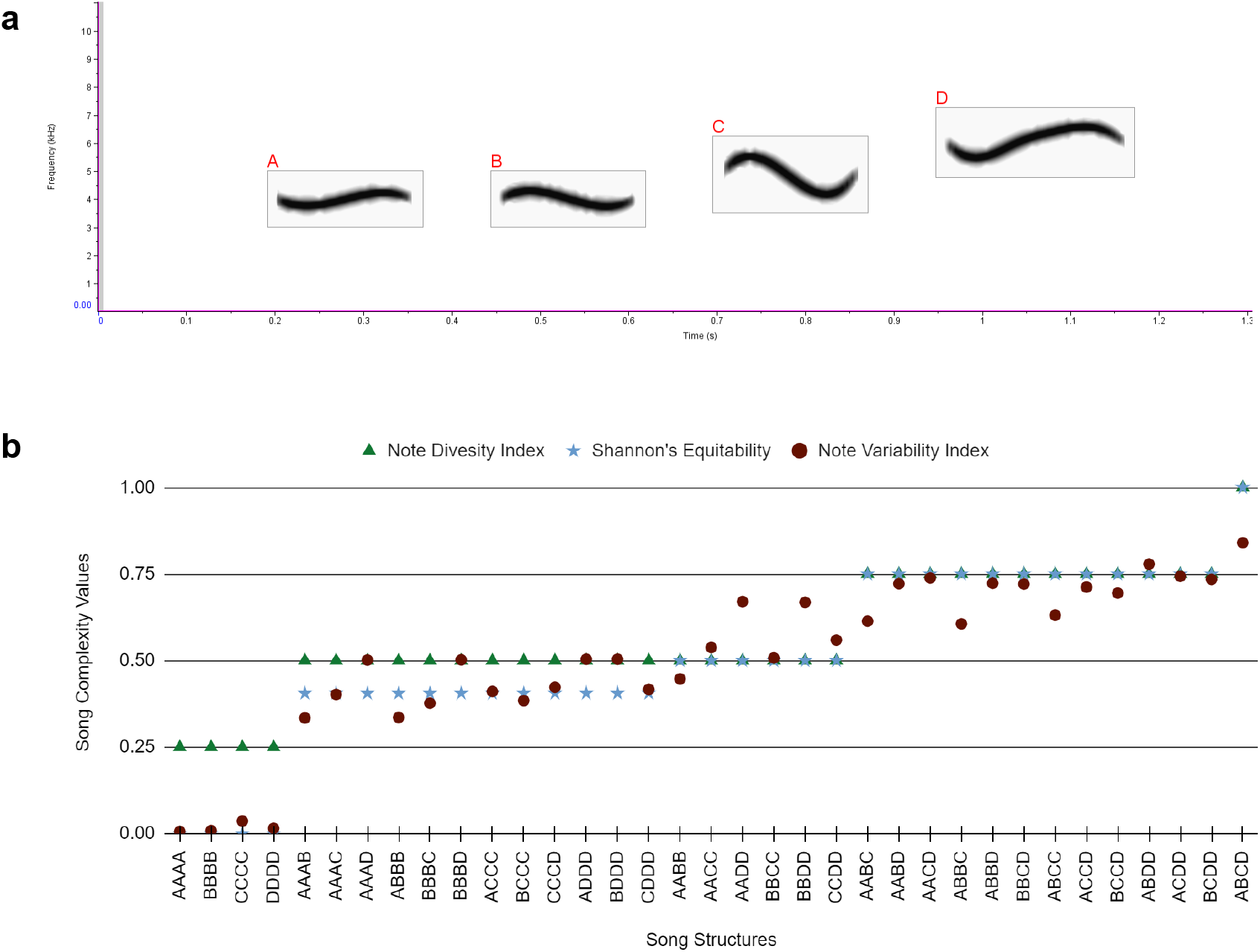
Synthetic songs are made of all possible combinations of four notes used to validate the complexity measures. (a) Spectrogram of four notes chosen with the note type labels - taken from White-bellied Sholakili song. (b)Note Diversity Index, Shannon’s Equitability, and Note Variability Index values for synthetic songs of 3 and 4 notes with all possible combinations of the four notes.

The output value (Formula 1) of NVI ∈ (0,1), where higher values represent high complexity and lower values represent low complexity.

#### Implementing NVI

Although the correlation coefficient (C_Δt_) can be arrived at using various methods (Pearson’s Correlation, Spearman’s Correlation, Kendall Correlation, Dynamic Time Warping of frequency contours, using defined spectral parameters), in this study we calculated C_Δt_ using the Spectrogram Cross-Correlation (SPCC) implemented as the ‘Correlator’ feature in the software Raven Pro 1.5. SPCC uses pixel-to-pixel matching between the spectrograms of the .wav files of each note (Charif et al., 2010). Correlation coefficient data for all notes (of each song) in this study were obtained through Raven Pro ‘Batch Correlator’ where the data using the same spectral parameters with Hann spectrograms window size of 512, an overlap of 50%, and a DFT size of 512.

### 2.1 Synthetic songs to compare Note Variability Index and other song complexity indices

We used the ‘sim_songs()’ function from the warbleR package (Araya-Salas & Smith-Vidaurre, 2017) to generate synthetic notes. For simplicity, we used two-steps (segments during Brownian motion) for notes within the frequency bounds of 4kHz to 6kHz. These notes consist of two very similar (relatively lower frequency) notes, one higher frequency note, and one between these (Fig.2a). We then calculated correlation coefficients of the notes using Spectrogram Cross-Correlation (SPCC) for each pair of these notes (Fig.2b), and for each song, we measured complexity using the NVI score along with two widely used measures, the ratio of note types and note count(N_T_ / N_C_) (denoted as Note Diversity Index) and Shannon’s Equitability 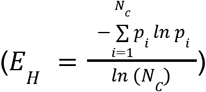 where N_T_ represents manually defined note types, and p_i_ is the stationary probabilities of each note in the song. We analyzed different structural groups of the synthetic songs to check for the advantages of NVI based on the inter-note variability (Fig.3c). We also compared these measures with the Functional Diversity measures (FRic, FEve, and FDiv) following its recent use to describe song complexity (Zsebők et al., 2021). (Supp. Fig 2)

**Fig. 2.**
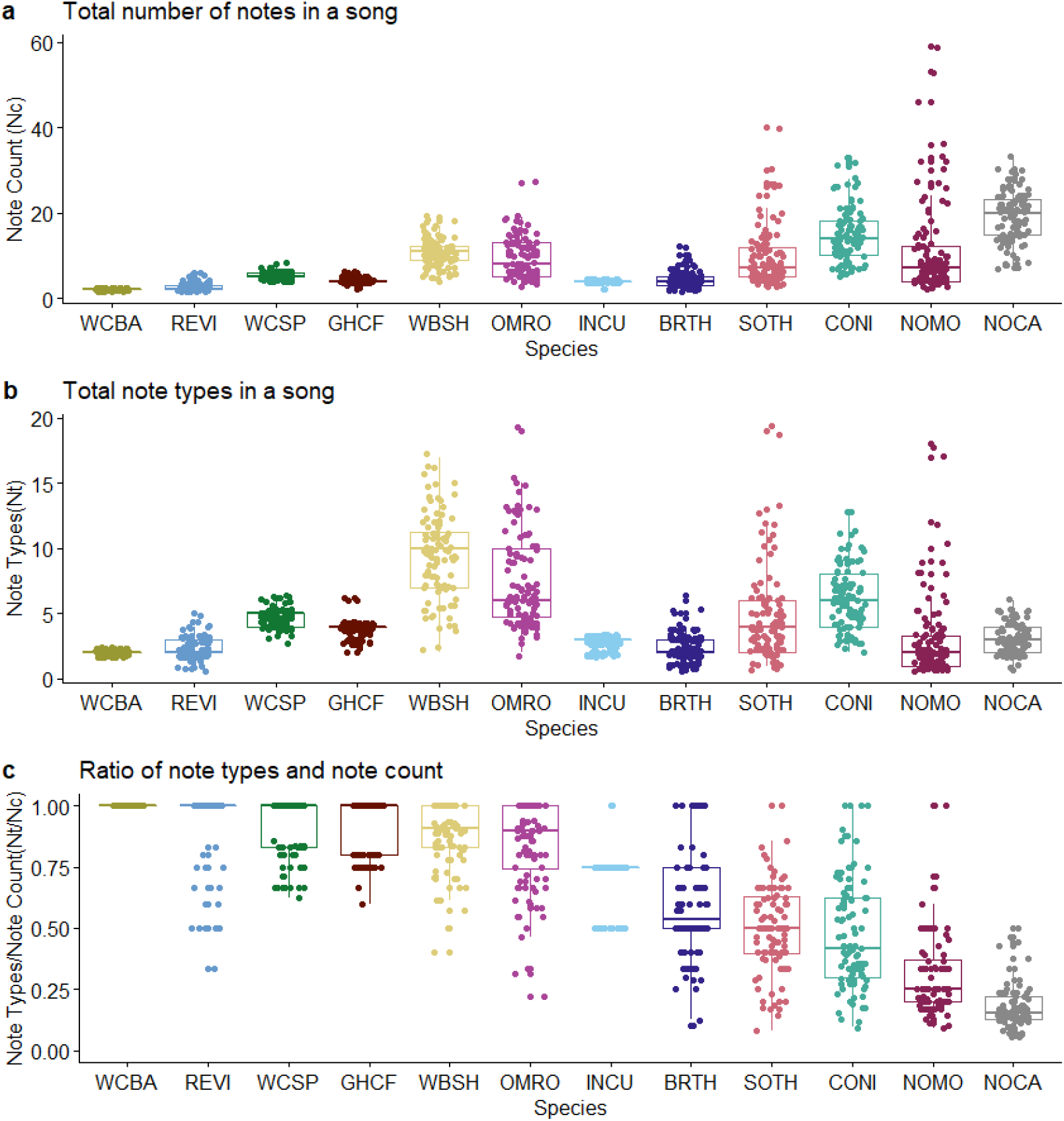
Comparison of different song parameters for songs of 12 species- WCBA- White-cheeked Barbet, REVI- Red-eyed Vireo, WCSP- White-crowned Sparrow, GHCF- Grey-headed Canary-Flycatcher, WBSH- White-bellied Sholakili, OMRO- Oriental Magpie-Robin, INCU- Indian Cuckoo, BRTH- Brown Thrasher, SOTH- Song Thrush, CONI- Common Nightingale, NOMO- Northern Mockingbird, NOCA- Northern Cardinal. (a)The total number of notes within a song(N_C_). (b) The number of manually classified note types within a song(N_T_). (c) Note Diversity Index(N_T_/N_C_).

**Fig. 3.**
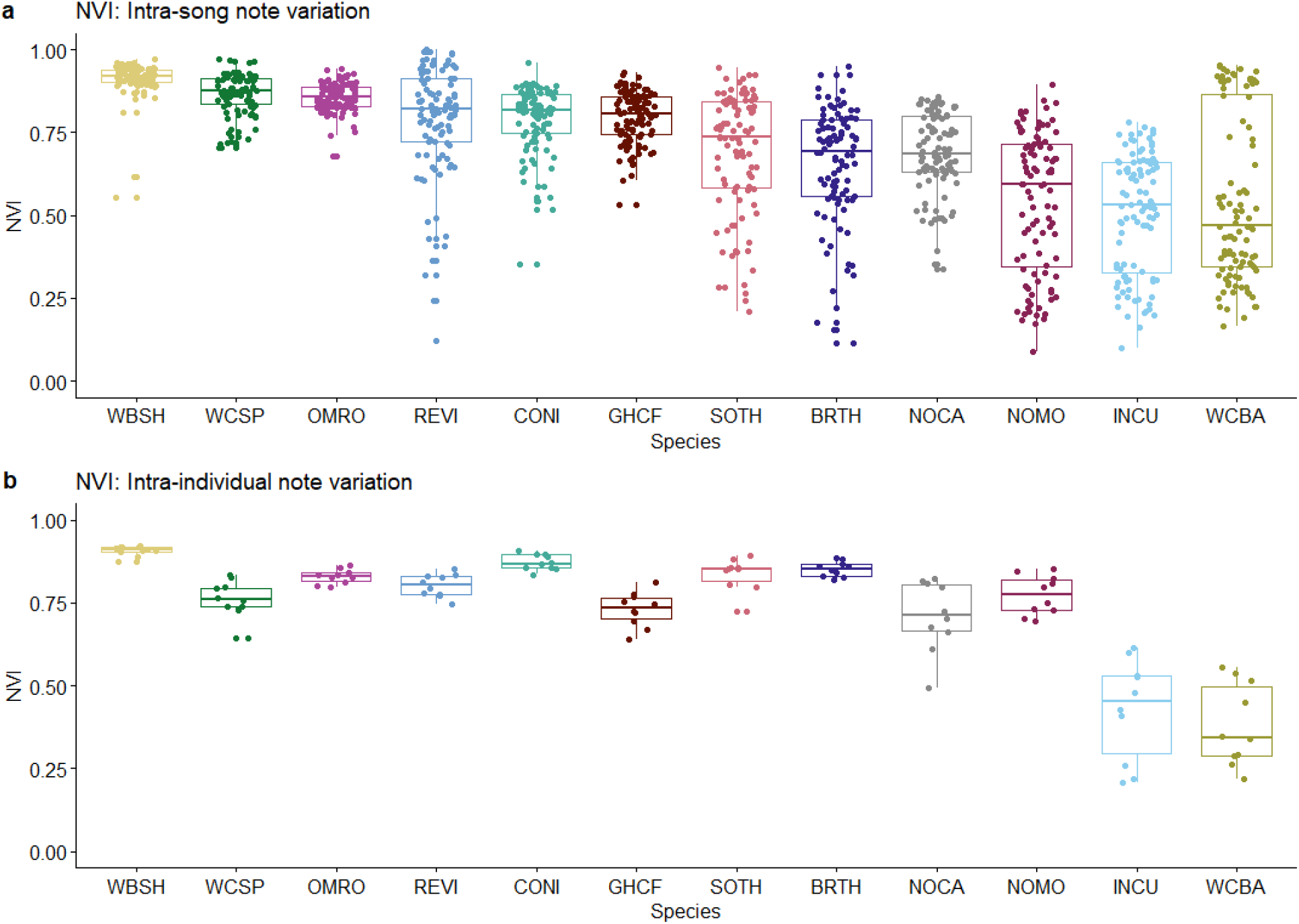
The Note Variability Index values for the 12 species- WBSH- White-bellied Sholakili, WCSP- White-crowned Sparrow, OMRO- Oriental Magpie-Robin, REVI- Red-eyed Vireo, CONI- Common Nightingale, GHCF- Grey-headed Canary-Flycatcher, SOTH- Song Thrush, BRTH- Brown Thrasher, NOCA- Northern Cardinal, NOMO- Northern Mockingbird, INCU- Indian Cuckoo, WCBA- White-cheeked Barbet. (a) Variation of all notes within a song- each dot represents a song. (b) Variation of notes across ten songs of an individual- each dot represents an individual

### 2.2 Note Variability Index for real birdsongs

To test measures of song complexity using NVI with real-world examples of song diversity, we used natural bird songs of twelve species occurring in different parts of the world. Based on existing literature, expert advice, and availability of acoustic data, we shortlisted six bird species that are believed to have complex songs which included the Brown Thrasher *Toxostoma rufum* (Boughey & Thompson, 1981), Common Nightingale *Luscinia megarhynchos (‘The top 10 British birdsongs’, n.d.; Weiss, Hultsch, Adam, Scharff, & Kipper, 2014*), Northern Mockingbird *Mimus polyglottos* (Derrickson & Breitwisch, 1992; Kroodsma, 2015), Song Thrush *Turdus philomelos (‘The top 10 British birdsongs’, n.d.*), White-bellied Sholakili *Sholicola albiventris* (Purushotham & Robin, 2016), and the White-crowned Sparrow *Zonotrichia leucophrys* (‘The Development of Birdsong’, n.d.; Nelson & Poesel, 2007). In addition, we chose six more species that are considered to have comparatively simpler songs which included -the Grey-headed Canary-Flycatcher *Culicicapa ceylonensis*, Indian Cuckoo *Cuculus micropterus*, Northern Cardinal *Cardinalis cardinalis*, Oriental-Magpie Robin *Copsychus saularis*, Red-eyed Vireo *Vireo olivaceus*, and White-cheeked Barbet *Megalaima viridis*. We obtained the song recordings of 11 species from the Macaulay Library (Cornell University), and we used the songs recorded by our lab for the White-bellied Sholakili. We collected songs of 10 individuals per species, assuming each separate recording to be from different individual birds, and chose ten songs per individual/recording for the analysis. This resulted in a total dataset of 100 songs per species.

The recordings and songs were selected based on the recording quality and the signal-to-noise ratio. All songs were manually annotated by one person (SS) in Raven Pro 1.5 (Charif et al., 2010). We counted the number of notes (N_C_) and types of notes (N_T_) and used these measures to calculate the Note Diversity Index (N_T_ / N_C_).

For NVI, we used species-specific frequency bandwidth cut-offs for the SPCC, which were based on the maximum and minimum frequencies for the fundamental notes of their vocalizations. We then calculated the NVI values for each song. As described in section 2.1, we normalized the immediate note variety represented by this NVI value with the total number of notes across ten songs to get a single song complexity value for each individual. The summary workflow is described in Appendix 1.

## Results

### 3.1 Comparison of complexity indices using synthetic songs

First, we compared two previously proposed complexity indices-Note Diversity Index (NDI) and Shannon’s Equitability (EH) with NVI using synthetic songs to enumerate the advantages and limitations of these methods. Fig. 1-b shows the complexity scores of synthetic songs generated from all possible combinations of three notes and four notes using these three complexity measures. The 55 possible combinations are divided into eight structural groups based on the number and repetition of notes. This showed differences in the complexities of songs within the same structural groups only for the NVI (Fig 3b). Due to the higher association within notes A and B, the NVI values for the synthetic songs with A and B co-occurring are relatively lower for the particular structural group. On the other hand, as the individual identities are lost in the case of Note Diversity Index and Shannon’s Equitability, all the combinations of the same structural groups show the same value for complexity.

### 3.2 Note types, note count, and Note Diversity Index for real bird songs

On comparing the complexity scores of different metrics on natural birdsongs, we saw that Northern Cardinal and Common Nightingale have longer songs. White-cheeked Barbet and Red-eyed Vireo have very short songs (Fig. 2-a). The values for note types(Fig. 2-b) are different as White-bellied Sholakili, Common Nightingale, and Oriental Magpie-Robin have relatively more note types within a song. Note Diversity Index suggests that Northern Cardinal, Northern Mockingbird, Common Nightingale, and Song Thrush have relatively less diversity. On the other hand, songs of other species show high NDI for most of the songs(Fig. 2-c).

### 3.3 Validating Note Variability Index using real bird songs

The Note Variability Index (NVI) was calculated to estimate note diversity within and across the songs of a species. NVI values for each species appear to be different from NDI. Notably, along with White-bellied Sholakili, White-crowned Sparrow, Oriental Magpie-Robin, and Red-eyed Vireo, the Common Nightingale, and Grey-headed Canary-Flycatcher show higher note variation. On the other hand, White-cheeked Barbet and Indian Cuckoo have low NVI values while they also have a higher N_T_/N_C_ ratio (Fig. 3-a).

To understand the overall note variety across songs, we calculated the NVI values across notes of ten songs without having manually defined song segment boundaries. The overall note variety for Song Thrush, Brown Thrasher, Northern Cardinal, and Northern Mockingbird is relatively higher than that of the immediate variety. And the overall variety for Indian Cuckoo and White-cheeked Barbet drops down compared to the immediate variety (Fig. 3-b).

## Discussion

In this study, we compared common song complexity measures with the newly developed Note Variability Index (NVI) measure. We found that NVI corresponds well with several song features that relate to the note diversity within a song. More importantly, it considers the spectral differences within notes while quantifying complexity which was not possible with previously used song complexity measures.

### Comparison with other song complexity measures

Most of the previously used complexity measures need the notes in a song to be classified into distinct categories, which could be difficult or inaccurate for birds with a highly variable vocal repertoire. Manual annotation of notes based on visual and audible differences can vary from person to person, and also, this process is very time-consuming. Certain notes with similar spectral features can be classified into the same or different note types based on personal opinions. For the notes with differences in spectral shape and being in the same frequency bandwidth, it is very difficult to classify them into separate categories (Fig. 4).

**Fig. 4.**
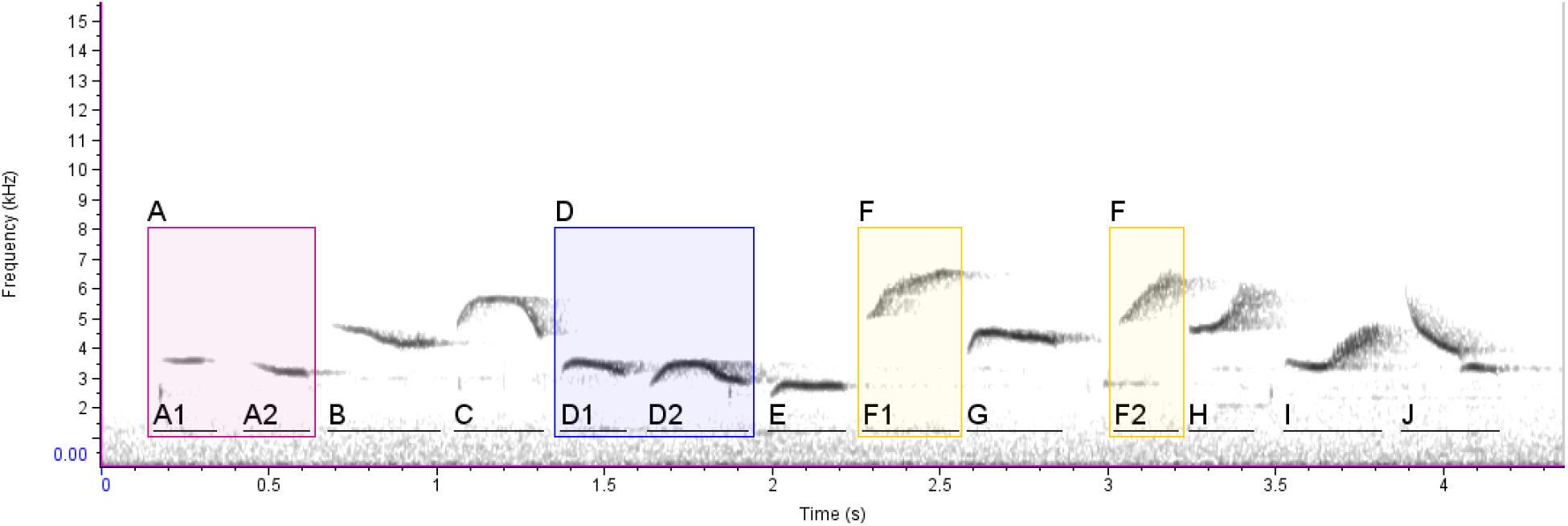
Spectrogram of White-bellied Sholakili songs with notes manually annotated into distinct classes-notes A1 and A2, D1 and D2, and F1 and F2 may or may not be classified as same notes A, D, and F respectively based on one’s perspective while doing manual annotation.

Though some studies have used specific algorithms to classify the notes into distinct categories, these algorithms’ efficiency and precision are uncertain (Pourhomayoun et al., 2013; Mcloughlin, Stewart, & McElligott, 2019; Rao, Montgomery, Garg, & Charleston, 2020).

### Limitations of functional diversity indices

Functional richness, evenness, and divergence used in Zsebők et al. (2021) can distinguish the note types by evaluating the spectral parameters. However, there may be limitations in using this to assess birdsong complexity. Acoustic parameters used for this measure are not enough to differentiate between the notes with similar bounds but with varying spectral shape (Supp. Fig. 2). Hence complexity values from Functional diversity indices may only work with songs with certain acoustic features - when spectral (note) shapes do not vary.

### Limitations of Note Diversity Index & Shannon’s Equitability

As Note Diversity Index and Shannon’s equitability don’t consider the extent of acoustic variation within notes, the resultant values may not interpret the real complexity of the songs. Also, the need for manual classification of notes into distinct categories makes these methods difficult and computationally slower. Notes within similar frequency bandwidths show higher association and result in less complexity (e.g., synthetic song Fig. 1-a, Notes C and D). For songs having similar structures, the NDI and SE give the same value of complexity as the notes are just considered as distinct units, and the extent of variation is not considered (e.g., synthetic song Fig. 1-b songs from ‘AABB’ to ‘CCDD’). As NVI is based on the spectral structure of individual notes, it can distinguish such songs.

### Immediate and Overall note variety

NVI also showed clear differences in immediate and overall note diversity for certain species. As NDI does not consider spectral parameters but bins notes into distinct categories, and only in cases where the manually classified notes are also spectrally divergent is there a consistency between NDI and NVI of immediate note variety (White-bellied Sholakili, Oriental Magpie-Robin, Red-eyed Vireo, and White-crowned Sparrow). In other cases, NDI may over-represent diversity (White-cheeked Barbet with only two marginally different notes has high NDI) or under-represent diversity (Northern Cardinal songs have notes across a larger frequency bandwidth but are repetitive and result in low NDI).

Some birds (Common Nightingale, Song Thrush, Brown Thrasher, and Northern Mockingbird) showed low immediate variety NVI values but high overall variety suggesting high turnover of notes across songs. On the other hand, other species (Indian Cuckoo and White-cheeked Barbet) had an opposite pattern suggesting a repetition of the same group of notes in every song.

### Interpreting levels of complexity with Note Variability Index

As NVI is calculated using the inverse of correlation coefficients, the magnitude of the final values may not be directly usable as a percentage of complexity. We suggest using Cohen’s standard to understand the association between correlation coefficients for evaluating the significance of the NVI value. NVI scores from 0.71 to 0.90 represent a higher complexity, NVI scores between 0.51 and 0.70 represent a medium complexity, and those of 0.50 and less represent a low complexity in terms of note diversity (Cohen, Cohen, West, & Aiken, 2013). These suggestions seem appropriate from the species analyzed here.

### Alternate methods for Note Variability Index computation

This study used the Spectrogram Cross-Correlation tool available in Raven Pro 1.5 to calculate NVI as it can be used efficiently for notes with variable acoustic structures. NVI can also be calculated with other tools that can quantify similarity between notes. The other classification methods using spectrogram correlations include Pearson’s Correlation, Spearman’s Rank Correlation, and Kendall’s Rank Correlation (Hauke & Kossowski, 2011), and these may be used for specific cases. Dynamic Time Warping of frequency contours (Daniel Meliza, Keen, & Rubenstein, 2013; Kaewtip, Alwan, O’Reilly, & Taylor, 2016), Soundpoints (Taft, 2011), features based classification (Greig & Webster, 2013), Hidden Markov Models (HMMs) (Kaewtip et al., 2016) and several deep learning methods like Convolution Neural Networks (CNNs) (Narasimhan, Fern, & Raich, 2017) are also widely used to classify acoustic signals. Based on the type of notes in a song, in some cases using some of these methods to calculate NVI can give more robust complexity values. Although we assessed several of these possibilities, we decided to use SPCC as implemented in RAVEN for its ease-of-use and wide applicability across species.

### Advantages of Note Variability Index

Based on the comparison with conventional song complexity measures, only NVI considers the spectral features of each note and calculates the complexity score to give higher values for songs with notes having variation in the spectral parameters and the shape. As the NVI does not require manual or automated classification of notes into distinct classes, the process is much faster and unbiased compared to the conventional methods. If precise automated detection of notes is possible, the NVI calculation will take even less time. The NVI can be easily implemented for any type of bird song and can be also used to calculate the complexity of any animal vocalization.

### Limitations of Note Variability Index

As NVI is focused on deriving complexity in terms of note diversity, it doesn’t consider stationary probabilities and order of notes which can be an important factor in certain studies. The Spectrogram Cross-Correlation used to compare notes while calculating the NVI scores may not be precise for unclear or noisy recordings. As the amount of noise increases-Signal-Noise Ratio (SNR) decreases, the correlation between two notes increases, eventually giving low NVI scores than expected (Sup materials). For recordings with some amount of background noise, various techniques like denoising (Brown, Garg, & Montgomery, 2018; Xie, Colonna, & Zhang, 2020) can be used before implementing NVI. Though denoising methods can improve the efficacy of NVI, it is recommended to use good quality focal recordings of the target species.

The Note Variability Index provides a better estimation for note diversity compared to that of the conventional measures. For cultural song evolution studies including song variation across species, across time-span, and across populations, the NVI can be easily implemented for large sets of data. Also, as NVI can be expanded for measuring complexity of any animal vocalizations, this method can be very helpful and a fast way to understand the complexity of animal vocalizations.

## Acknowledgments

We acknowledge receipt of media from The Macaulay Library at the Cornell Lab of Ornithology and thank Matt Medler of the Macaulay Library for assistance in listing complex bird songs and obtaining the acoustic data. We thank Michael Pitzrick, Russ Charif, and Laurel Symes of the Center for Conservation Bioacoustics (CCB) for their constructive feedback while developing the new song complexity index, especially during an internship visit of SS to CCB. We thank the members of the two Ecology Labs at IISER Tirupati, in particular Harikrishnan C P, Isha Bopardikar, and Vijay Ramesh, for their feedback and support. This study forms part of the Master’s dissertation of SS as part of his BSMS course. We acknowledge the support of IISER Tirupati.

## Authors Contributions

S.S., C.A., V.J., and V.V.R.. contributed to the conceptualization and implementation of the study; S.S., C.A., and V.V.R. developed the new complexity method; S.S. acquired and analyzed the bird song data and led the manuscript writing process with revisions provided by all authors.

## Data Availability

1. R code for calculation of NVI from SPCC output- https://github.com/suyash-sawant/Birdsong_Complexity/blob/main/NVI_calculation.R

