## Supplemental Information for "Spectrogram cross-correlation can be used to measure the complexity of bird vocalizations"

### Supporting Information

| Complexity measure | Studies using the measure | Diversity of Notes | Rarity of Notes | Order of Notes | Spectral parameters of Notes | Spectral shpae of Notes | Manual Classification of Notes needed? |
|---|---|---|---|---|---|---|---|
| Shannon's Equitability | Kershenbaum, 2014 | ✘ | ✓ | ✘ | ✘ | ✘ | ✓ |
| Repeat Distribution | Kershenbaum & Garland, 2015 | ✘ | ✓ | ✓ | ✘ | ✘ | ✓ |
| Mutual Information | Kershenbaum & Garland, 2015 | ✘ | ✓ | ✓ | ✘ | ✘ | ✓ |
| Markov Models | Katahiraet al., 2013; Kershenbaum & Garland, 2015 | ✓ | ✘ | ✓ | ✘ | ✘ | ✓ |
| Lempel-Ziv Complexity Measure | Kershenbaum, 2014; Kershenbaum & Garland, 2015 | ✘ | ✘ | ✓ | ✘ | ✘ | ✓ |
| Levenstein Distances | Kershenbaum, 2014; Kershenbaum & Garland, 2015 | ✘ | ✘ | ✓ | ✘ | ✘ | ✓ |
| Entropy Rate | Kershenbaum, 2014; Kershenbaum & Garland, 2015 | ✓ | ✓ | ✓ | ✘ | ✘ | ✓ |
| Song Directed Networks | Sasahara et al., 2012 | ✘ | ✘ | ✓ | ✘ | ✘ | ✓ |
| Song Undirected Networks | Sasahara et al., 2013 | ✘ | ✘ | ✓ | ✘ | ✘ | ✓ |
| Repertoire Size | László Z. Garamszegi et al., 2005; Zeng et al., 2007; Petrusková et al., 2016; Benedict & Najar, 2019 | ✓ | ✘ | ✘ | ✘ | ✘ | ✓ |
| Note Type / Note Count | Spenceret al., 2003 | ✓ | ✘ | ✘ | ✘ | ✘ | ✓ |
| Independent Song Features | Benedict & Najar, 2019 | ✓/✘ | ✘ | ✘ | ✓/✘ | ✘ | ✓/✘ |
| PCA of Song features | Mason, Shultz, & Burns, 2014 | ✓/✘ | ✘ | ✘ | ✓/✘ | ✘ | ✓/✘ |
| Functional Diversity measures | Zsebők et al., 2021 | ✓ | ✘ | ✘ | ✓ | ✘ | ✘ |
| Note Variability Index | | ✓ | ✘ | ✘ | ✓ | ✓ | ✘ |

Supplementary Figure 1: Advantages and limitations of previously published song complexity measures compared to that of NVI

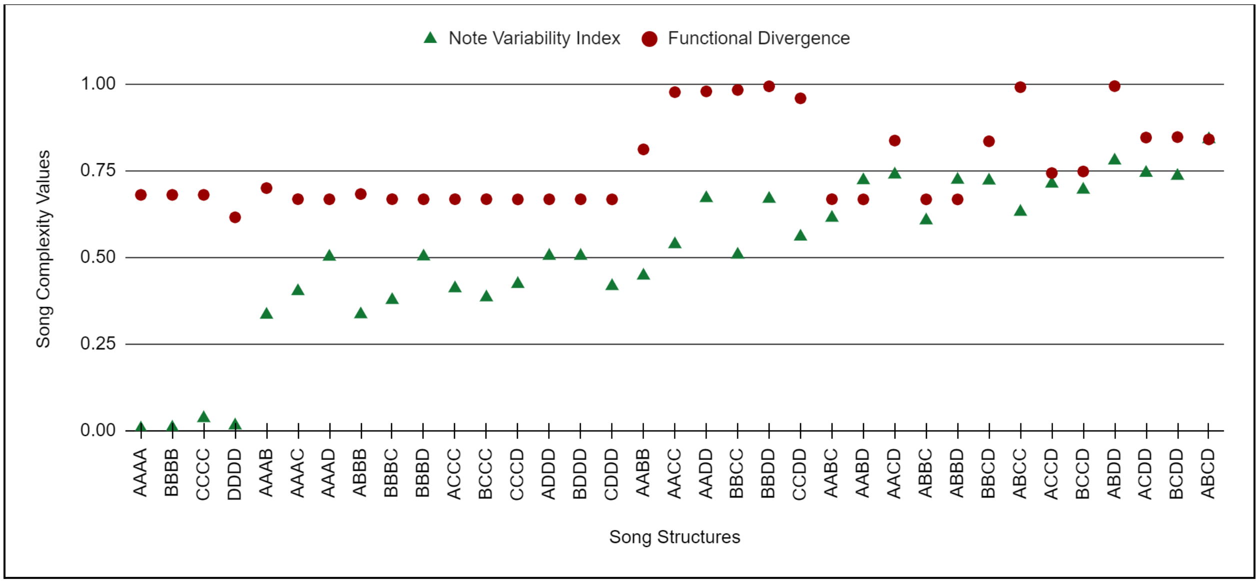


Supplementary Figure 2: Comparison of Note Variability Index and Functional Divergence for synthetic songs

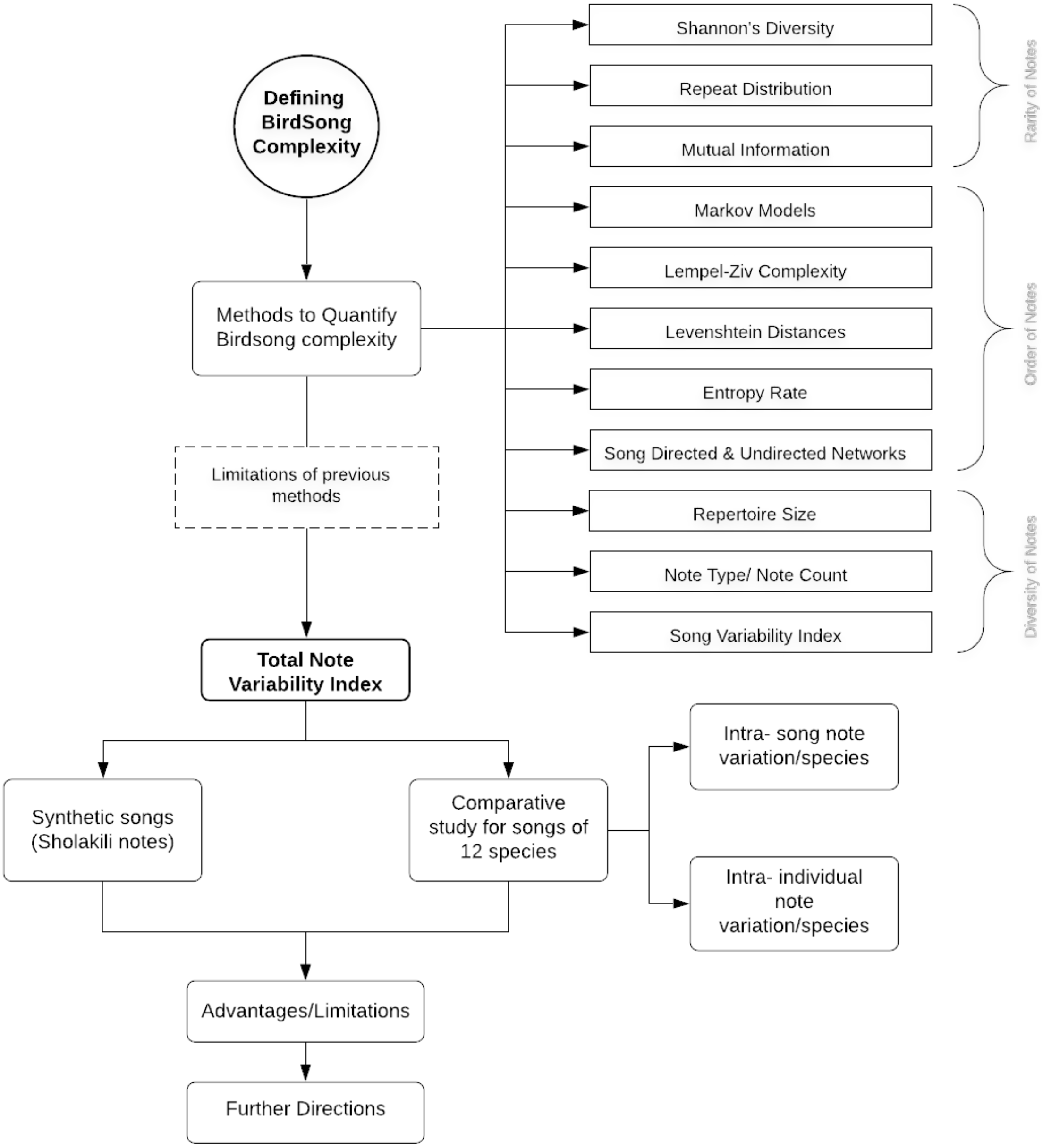


**Appendix. 1.** Flowchart showing the workflow of this study

| New File Name | Catalog Number |
|---|---|
| BRTH_01_10 | ML 11182 |
| BRTH_02_10 | ML 50243 |
| BRTH_03_10 | ML 52875 |
| BRTH_04_10 | ML 67309 |
| BRTH_05_10 | ML 79394 |
| BRTH_06_10 | ML 84762 |
| BRTH_07_10 | ML 94278 |
| BRTH_08_10 | ML 98812 |
| BRTH_09_10 | ML 100755 |
| BRTH_10_10 | ML 107293 |
| CONI_01_10 | ML 11276 |
| CONI_02_10 | ML 11280 |
| CONI_03_10 | ML 11281 |
| CONI_04_10 | ML 23656 |
| CONI_05_10 | ML 36169 |
| CONI_11_10 | ML 86188 |
| CONI_07_10 | ML 36187 |
| CONI_12_10 | ML 71614 |
| CONI_13_10 | ML 84565 |
| CONI_10_10 | ML 71609 |
| GHCF_01_10 | ML 173508 |
| GHCF_11_10 | ML 183373 |
| GHCF_03_10 | ML 184655 |
| GHCF_04_10 | ML 183144241 |
| GHCF_05_10 | ML 197734 |
| GHCF_06_10 | ML 202068 |
| GHCF_07_10 | ML 182961 |
| GHCF_08_08 | ML 175170 |
| GHCF_09_02 | ML 168546 |

| New File Name | Catalog Number |
|---|---|
| NOCA_01_10 | ML 209312 |
| NOCA_02_10 | ML 191165 |
| NOCA_03_10 | ML 176241 |
| NOCA_04_10 | ML 134151 |
| NOCA_05_10 | ML 130905 |
| NOCA_06_10 | ML 107306 |
| NOCA_07_10 | ML 98872 |
| NOCA_08_10 | ML 94248 |
| NOCA_09_10 | ML 84683 |
| NOCA_10_10 | ML 77283 |
| NOMO_01_10 | ML 534555 |
| NOMO_02_10 | ML 249820 |
| NOMO_03_10 | ML 239240 |
| NOMO_04_10 | ML 237660 |
| NOMO_05_10 | ML 210543 |
| NOMO_11_10 | ML 23402 |
| NOMO_07_10 | ML 100752 |
| NOMO_08_10 | ML 166628 |
| NOMO_09_10 | ML 133234 |
| NOMO_10_10 | ML 118628 |
| OMRO_01_10 | ML 52951 |
| OMRO_02_10 | ML 177229 |
| OMRO_11_10 | ML 70503 |
| OMRO_12_10 | ML 284631 |
| OMRO_05_10 | ML 184615 |
| OMRO_06_10 | ML 198994 |
| OMRO_07_10 | ML 212082 |
| OMRO_08_10 | ML 215056 |
| OMRO_13_10 | ML 11197 |

| New File Name | Catalog Number |
|---|---|
| SOTH_01_10 | ML 14083 |
| SOTH_02_10 | ML 36174 |
| SOTH_11_10 | ML 249163 |
| SOTH_04_10 | ML 77510 |
| SOTH_05_10 | ML 162845 |
| SOTH_06_10 | ML 204480 |
| SOTH_12_10 | ML 14085 |
| SOTH_08_10 | ML 231137 |
| SOTH_09_10 | ML 231157 |
| SOTH_10_10 | ML 236521 |
| WBSH_01_10 | RV SHAL_001 |
| WBSH_02_10 | RV SHAL_002 |
| WBSH_03_10 | RV SHAL_003 |
| WBSH_04_10 | RV SHAL_004 |
| WBSH_05_10 | RV SHAL_005 |
| WBSH_06_16 | RV SHAL_006 |
| WBSH_07_10 | RV SHAL_007 |
| WBSH_08_10 | RV SHAL_008 |
| WBSH_09_10 | RV SHAL_009 |
| WBSH_10_15 | RV SHAL_010 |
| WCBA_01_10 | ML 237520 |
| WCBA_02_10 | ML 213245 |
| WCBA_11_10 | ML 166138 |
| WCBA_12_10 | ML 24774 |
| WCBA_05_10 | ML 283231 |
| WCBA_06_10 | ML 283156 |
| WCBA_13_10 | ML 559 |
| WCBA_14_10 | ML 569 |
| WCBA_09_10 | ML 283044 |

| | | | | | |
|---|---|---|---|---|---|
| GHCF_10_05 | ML 175272 | OMRO_10_10 | ML 36417 | WCBA_15_10 | ML 726 |
| INCU_01_10 | ML 178382 | REVI_01_10 | ML 11830 | WCSP_01_10 | ML 16627 |
| INCU_02_10 | ML 36293 | REVI_02_10 | ML 11867 | WCSP_02_10 | ML 16641 |
| INCU_03_10 | ML 175427 | REVI_03_10 | ML 38556 | WCSP_03_10 | ML 16676 |
| INCU_04_10 | ML 176989 | REVI_04_10 | ML 55513 | WCSP_04_10 | ML 22966 |
| INCU_05_10 | ML 202521 | REVI_05_10 | ML 63933 | WCSP_05_10 | ML 42260 |
| INCU_06_10 | ML 146150 | REVI_06_10 | ML 67811 | WCSP_06_10 | ML 49958 |
| INCU_07_10 | ML 68711 | REVI_07_10 | ML 73894 | WCSP_07_10 | ML 49990 |
| INCU_11_10 | ML 169312 | REVI_08_10 | ML 79425 | WCSP_08_10 | ML 50020 |
| INCU_09_10 | ML 212077 | REVI_09_10 | ML 84847 | WCSP_09_10 | ML 50105 |
| INCU_10_10 | ML 180679 | REVI_10_10 | ML 105271 | WCSP_10_10 | ML 66714 |

**Appendix. 2.** Availability of data used for analyzing natural birdsongs
